# Simulations of higher-order protein assemblies using a fuzzy framework

**DOI:** 10.1101/248062

**Authors:** B. Tüű-Szabó, L. Kóczy, M. Fuxreiter

## Abstract

Spatiotemporal regulation of the biochemical information is often linked to supramolecular organizations proteins and nucleic acids, which generate membraneless cellular organelles. Owing to difficulties in high-resolution structural studies, the driving forces of assembling these low-complexity polymers have yet to be elucidated. Polymer physics approaches captured the experimentally demonstrated critical role of binding element multivalency and highlighted the importance of linker solvation. Here we present a simulation method based on a fuzzy mathematical framework. This approach is suitable to handle the heterogeneity of interactions pattern generated by redundant binding motifs and the resulted multiplicity of conformational states. Using a hypothetical polymer, fuzzy simulations recapitulate the experimental observations on valency-dependence and are more efficient than the one-to-one binding model. Systematic studies on binding element affinity and linker dynamics demonstrate that these two factors present alternative scenarios to promote polymerization: stronger binding result in more ordered states, whereas increasing dynamics contributes to heterogeneity and a more favorable entropy of the assembly. We propose that the fuzzy framework could be employed to characterize/predict mutations leading to pathological aggregates.

## Introduction

Proteins can form a wide variety of assemblies, regarding composition, size, and dynamics. In addition to simple binary, ternary complexes and middle-size oligomers, proteins may also assemble into higher-order organizations. These supramolecular assemblies are implicated in different biological processes ranging from normal physiology to disease [1, 2]. For example, to minimize signaling noise for low-affinity effectors signaling complexes frequently increase local concentration of binding sites via higher-order protein assembly [3]. Recent discoveries revealed that supramolecular organizations of proteins and nucleic acids can generate functional cellular compartments [4, 5]. They appear at various points on the biological landscape and usually lack a membrane boundary [6]. Such membraneless organelles can serve as biomolecular storages upon stress, bioreactors to accelerate chemical reactions as well as signaling devices, which assembly/disassembly is regulated by a variety of pathways [7-9]. Although such cellular bodies, for example the nucleolus functioning in ribosomal RNA transcription were discovered long ago, their molecular basis and the underlying physical forces have remained largely enigmatic. Seminal works by the Brangwynne, Hyman, Parker labs revealed that these organelles are created by a process of liquid-liquid demixing, once the component concentration exceeds the saturation limit [10, 11]. This phenomenon, also termed as phase transition have been described by polymer physics [12]. In this sense, protein chains could be considered as biological polymers, which could be crosslinked to generate a large, system-spanning network.

Later studies revealed that proteins, which participate in this process have degenerate binding motifs and their sequences are often composed of tandem repeats of a few residues [13]. Owing to their information content, these sequences are also termed as low-complexity (LC) regions or domains. Multivalency is a critical component of phase transition, as it was demonstrated by the Rosen lab using engineered SH3 domains and interacting proline-rich motifs [14]. Another interesting feature is that proteins, which form membraneless organelles often possess long segments without a well-defined tertiary structure, termed as intrinsically disordered proteins/regions (IDPs/IDRs) [15]. The relationship between the dynamics of these disordered segments and the characteristics of the resulted membraneless organelles have yet to be elucidated.

Higher-order protein organizations exhibit a wide spectrum of states with distinct dynamics. Prions/amyloids are stabilized by β-zippers, resulting in static, solid-like inheritable entities [16]. Signalosomes, such as inflammasomes or necrosomes could resemble prion-like, stable structures [17] or be dynamic, for example the autophagosome [18]. Ribonucleoproteins (RNP) generate dynamic granules or liquid-like droplets [19]. Nuclear pore complexes (NPCs) are somewhat more stable and form hydrogels [20]. Intriguingly, the very same protein could be organized into higher-order states with distinct dynamics. Pathological mutations may induce conversion of liquid-like droplets to solid fibrils, for example in case of the Fus or hnRNPA protein, familial mutations of which appear in Amyothrophic lateral sclerosis (ALS) [1, 2]. In order to understand how such aberrant transitions occur, molecular factors determining the material state have to be determined.

Recently, a unified framework for higher-order structures has been proposed, which decomposed the material state to three factors [21]. First, low-affinity interacting elements, such as cation-pi, pi-pi, aromatic hydrogens bonds are required for a dynamical equilibrium with fast off-rates. Second, degenerate interactions between the binding elements increase the number of microstates and results in favorable entropy. Third, protein segments must preserve their conformational entropy to enable interactions in a vast number of arrangements. This phenomenon, which is referred to as fuzziness [22, 23], is a universal feature of all higher-order assemblies [21]. Indeed, NMR data indicates similar conformational heterogeneity of Fus in its free and bound states [19]. Modelling the effect of fuzziness however is challenging, owing to the large system size and entropy.

Both microscopic and macroscopic methods could be applied to gain insights into the driving forces of the assembly. In the microscopic, polymer physics treatment, intramolecular interactions within the individual polymers are exchanged for intermolecular interactions between the different chains to form reversible, system-spanning crosslinks [12]. This process could be described by the Flory–Huggins theory [24], which quantifies the critical concentration and explains the surface tension of the droplets. In a macroscopic approximation, phase separation is driven by solvation effects, when the condensed biomolecular environment provides a more favorable environment for protein chains than the solvent [25]. A simplified polymer physics approach has been applied to recapitulate the valency-dependence of phase transition of the artificial system containing SH3 domains and proline-rich motifs [14]. Despite the heavy assumptions of the model, such as the dimensionless molecules with no upper boundary for the concentration, linkers with infinite flexibility to enable any kinds of binding arrangements, and the lack of cooperativity between the different binding sites, these rule-based stochastic simulations provided results in agreement with the experimental observations as well as with theoretical estimates [14]. Recently, a more detailed Monte Carlo simulation on a coarse-grained lattice was applied on the same system [26]. This model enabled to study the properties of the linkers, using the effective solvation volume as a single parameter. This quantity, which was derived from allatom simulations, expressed the preference of the linkers for themselves or the solvent, based on which compact and extended chains could be distinguished. The results shed light on how different protein sequences affect phase transition, depending on their compaction and solvation properties [26, 27]. Neither of these techniques however, could account for the conformational properties (i.e. dynamics) of the linkers, which could be dramatically altered by pathological mutations.

We developed a novel approach, which exploits the intrinsic fuzziness of higher-order protein organizations [21]. Here fuzziness refers to the function-related structural heterogeneity of proteins, which is resulted by redundant/degenerate, often transient interactions [28, 29]. Such conformation and interaction heterogeneity can enable adaptive transitions in proteins [23]. The importance of fuzziness on regulated organization and activity of protein-protein or protein-nucleic acid assemblies has been demonstrated experimentally [30]. Fuzziness is also known as a mathematical concept, where the membership in given sets is described by a function, varying between [0,1] instead of a single [0 or 1] value. Fuzziness has been derived from the seminal work of Lotfi Zadeh [31], and has been implicated in the electronic control of ∼3000 artificially intelligent devices [32]. Mathematical fuzziness however, has not been applied to protein interactions.

Here we attempt to use the mathematical concept of fuzziness to describe phase transition of low-complexity protein sequences. In this model, a single binding site can simultaneously interact with multiple binding elements to different extents, in contrast to the previous approaches, where only one-to-one binding was considered. We demonstrate that incorporation of fuzzy relations between the degenerate interaction elements not only increases efficiency of the simulations, but also enables to account for other important factors such as local concentration or linker dynamics.

## Methods

### Model system

The model is a hypothetical biological polymer, which is composed of N residues. Each residue is characterized by two values: binding preference and dynamics. Both values vary in a [0,1] range. Binding preference > 0.3 designate residues involved in binding, dynamical values > 0.3 correspond to linker residues. Binding elements (α) are defined as a continuous stretch of at least 5 residues with binding preference > 0.3. Linkers (λ) are defined as all other segments connecting the binding elements. If the length of the linker is ≤ 3, the two connected binding elements are merged. These two characteristics (binding preference, dynamics) enable to consider binding elements with varying affinities as well as allow linkers to form transient contacts.

### Parameters

Binding preference of a binding element (α_i_) is obtained as an average of the residue binding preferences:

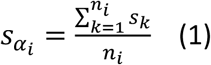

where *s*_*k*_ is the binding preference of residue *k* and the binding element contains *n*_*i*_ residues.

Affinity between two binding elements is defined as the average of the binding preferences [0,1]:

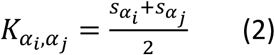

where *s*_*α_i_*_.and *s*_*α_j_*_ are the binding preferences of the interacting α_i_ and α_j_ elements.

As the fuzzy framework allows one binding element to interact with multiple other elements, the number of possible binding elements available for interaction needs to be determined. All binding elements within the volume defined by the neighboring linkers are considered:

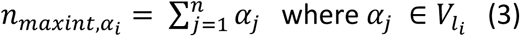

where *n_maxintαi_.* is the maximum number of interaction sites around the binding element α_i_. The available volume for α_i_ interactions is defined by the length of the longer neighboring linker (l_i_). In the default case, *V*_*l_i_*_. is a spherical volume with a radius of *l_i_*,:

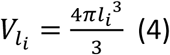

This volume could be re-scaled according to linker dynamics, which is defined as the average of the residue dynamics:

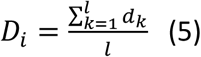

where *d*_*k*_ is the dynamical value of residue *k*, and *l*_*i*_ is the length of the linker.

If the linker dynamics (*D*_*i*_) is 1, all the available binding sites are considered within the volume as defined in eq. 4. If *D*_*i*_) < 1, the sphere radius is reduced proportionally to the linker dynamics:

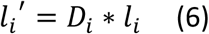

where *D*_*i*_ is the linker dynamics and *l*_*i*_ is the length of the linker. *l′*_*i*_ is used to obtain the volume by eq 4.

### Computed quantities

The association probability linearly depends on the binding affinity (*K*_*α_i_*_,_*α_j_*_), and reciprocally on the available binding sites (*n*_*maxint*_,*α*_*i*_)

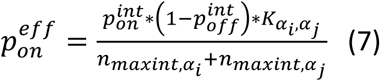

where the intrinsic association probability 
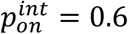
, the intrinsic dissociation probability 
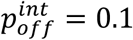
, similarly to the reference [14].

Once interactions are formed in the first step, we define an occupancy value for each binding elements [0,1]. It is calculated with an algebraic sum, which is the fuzzy union (s-norm) operator.

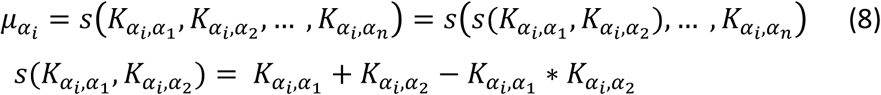

where *n* is the number of the binding elements in the polymer.

From the second step, the affinity of a given interaction between α_i_ and α_j_ also depends on the local concentration of the bound binding elements. The binding affinity between two elements must be weighted by the occupancies of the neighboring binding sites:

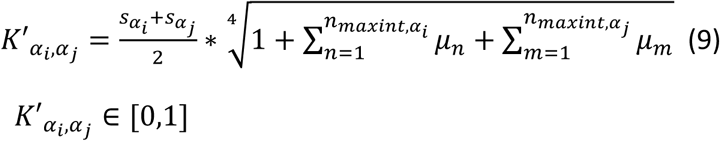

where *μ*_*n*_ and *μ*_*m*_ occupancies are summarized for all binding elements within the available volume for α_i_ and α_j_. If local concentration is taken into account from the second step, the modified affinities (*K′*_*α_*i*_*_,_*α_*j*_.*_) are used to determine the occupancies in eq. 8.

The association probability is also modified accordingly:

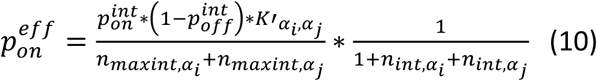

where *K′*_*α_*i*_*_,_*α_*j*_*_ is the modified binding affinity defined in eq. 9, *n*_*int*_,*α*_*i*_ and *n*_*int*_,*α*_*j*_. are the actual number of binding elements, which are bound to α_i_ and α_j_, respectively. The dissociation probability has an inverse relationship to the binding affinity:

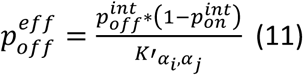

where the intrinsic association 
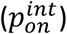
 and dissociation 
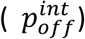
 probabilities are the same as in eq. 7, and *K′*_*α_i_*_,_*α_j_*_. is the modified binding affinity defined in eq. 9.

From the second step, the molecules could be present as individual chains, or chains organized into oligomers or larger polymers. Here we need to define the interaction capacity (‘freedom’) of the binding elements within the molecular assembly/polymer:

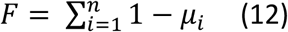

where (1 — *μ*_*i*_) is the available interaction capacity of a given α_i_ binding element, which is summarized for all binding elements in the polymer.

Finally, any molecule types in the system: individual molecules, oligomers or larger polymers can diffuse to another box. The probability of the diffusion is defined as:

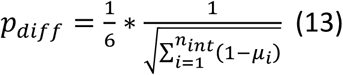

where the square root of *F* (eq. 12) is used in the denominator.

### Computational protocol

A periodic system is defined, which contains Z boxes, with dimension w. The system contains m molecules, each composed of N residues. Initially the m molecules are placed randomly in the cubes. The simulation protocol is based on the stochastic rule-based simulation in reference [14].

In the first simulation step the molecules can associate according to the probabilities, which are defined in eq 7. From the second iteration step, the molecules have three options: they can associate (*i*), dissociate (*ii*) and diffuse to another, randomly chosen neighboring cell (*iii*). Affinities of the interactions depend on the local concentration of the bound elements, therefore occupancies (eq. 8) and the interaction capacities (eq. 12) are determined in each step. Association probabilities (eq 10) and dissociation probabilities (eq 11) are proportional to the modified affinities, so both account for the local concentration of the available sites as well as their binding status. Diffusion is inversely proportional to the interaction capacity, so larger polymers have less chance to move to the neighboring box. Linker dynamics is taken into account through the volume, within which the available binding elements are computed.

## Results

The fuzzy models allowed heterogeneous interactions with multiple binding elements to different extents, in contrast to the non-fuzzy model, where contacts are established with a single, well-defined binding partner. All previous simulations of higher-order assemblies were based on one-to-one interactions [14, 26]. We performed simulations with fuzzy and the one-to-one (non-fuzzy) binding models and compared the probabilities of formation of large polymers, which were defined as interconnected m > 25 molecules. Using hypothetical molecules characterized by binding affinity and dynamics varying in [0,1] range, we studied the impact of valency on polymer formation. The model system contained one molecule type, the size of which has been systematically varied between 2 to 6 binding elements and linkers, the length of which were arbitrary defined as 7 and 10 residues, respectively (Figure 1). The system contained m molecules randomly positioned in Z boxes. Concentration was modulated by varying the size of the simulation box. The topologies and the parameters considered for interactions are shown in Figure 1. Three types of stochastic movements were performed, similarly to the reference [14]: i) association ii) dissociation iii) diffusion to a neighboring box. In addition to valency, we could monitor the impact affinity and local concentration of binding elements as well as linker dynamics on the formation of large polymers. The results are averaged for 10 parallel simulations (10000 steps) for each parameter combinations.

**Figure 1.**
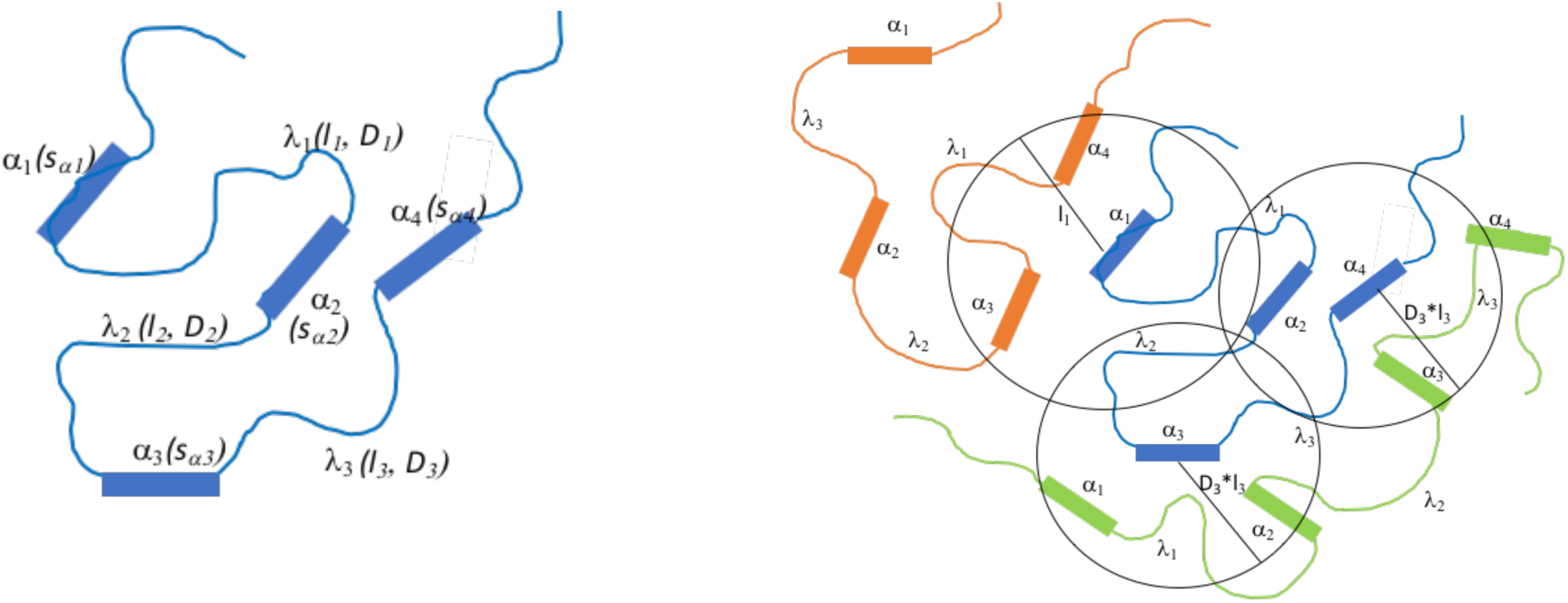
*Schematic representation of the model system with one (**left**) and three interacting molecules (**right**). sα_i_ is the binding preference of a binding element (α), computed as the average of the residue-based binding affinities (eq. 1). D_i_ is the dynamics of linker (λ) with length l_i_, obtained as the average of the residue-based dynamics values (eq. 5). Local concentration of the available binding sites is computed within a volume V_*l_i_*_., which is scaled by the linker dynamics (D_i_, eq. 6).*

Multi-valency is considered as the major driving force of phase transition [6, 14]. The impact of valency and concentration on the probability of formation of large polymers are shown in Figure 2. Concentration is given in arbitrary units, which decreases with increasing cube size. Both one-to-one and fuzzy simulations show strong dependence on the number of binding elements, recapitulating previous experimental observations [14].

**Figure 2.**
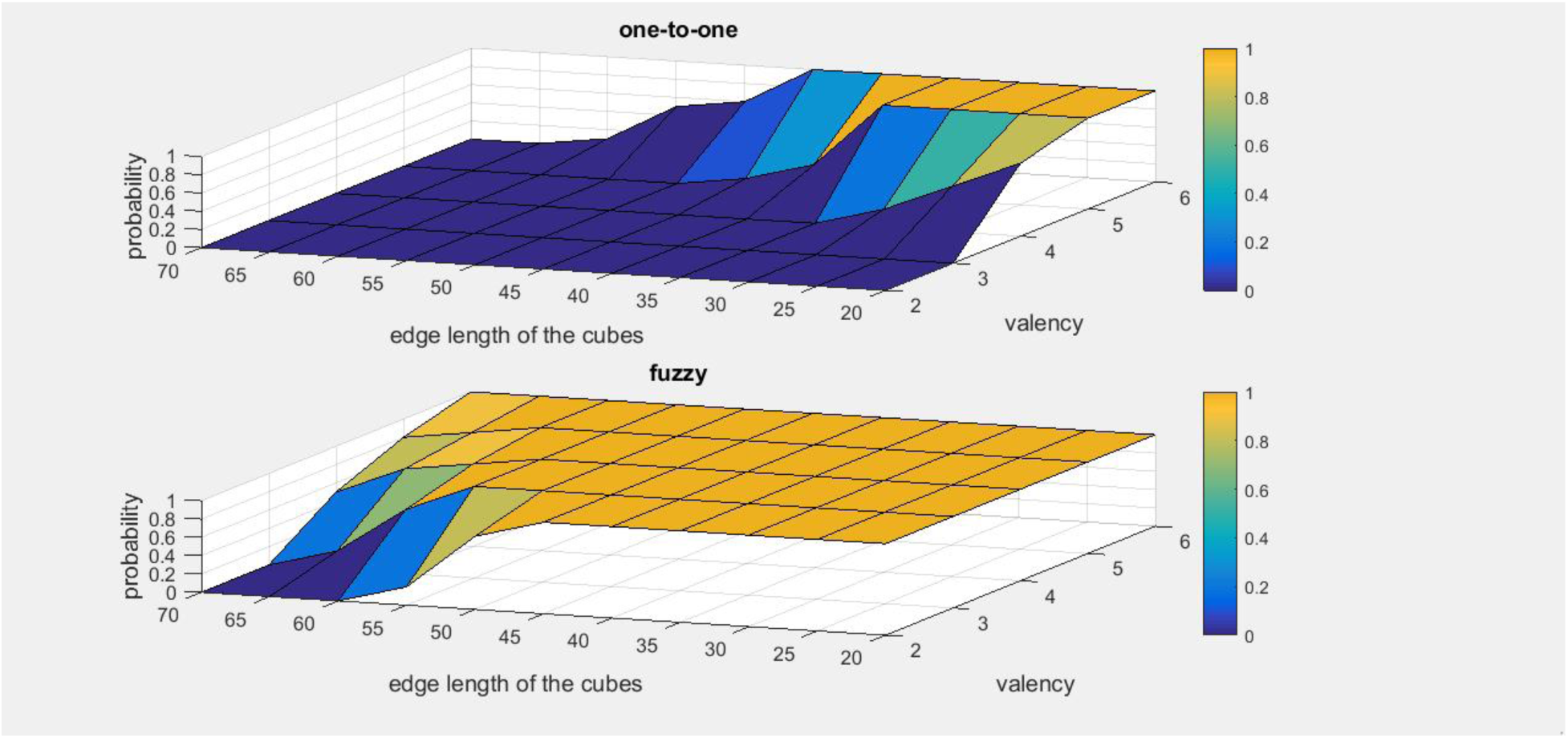
*Probability of large polymers as a function of valency and edge length in one-to-one (**upper**) and fuzzy (**lower**) binding models. Valency is defined as the number of binding elements. Concentration is given by the number of molecules/volume (L^3^). Binding element affinity=1.0, linker dynamics=1.0.*

Heterogeneous interactions with multiple partners enable polymer formation even at lower valency. Indeed, polymerization in fuzzy simulations occur almost at one order of magnitude lower concentration than in the one-to-one binding model (Figure 2). In simulations, which were conducted using lower binding element affinities (0.35) and linker dynamics (0.35) no higher-order oligomers (m > 25) were observed in the one-to-one simulations, while polymers were formed in case of > 4 binding elements in the fuzzy model (Figure 3). This illustrates that multi-valency influences polymerization via generating a large number of microstates and improved binding entropy.

**Figure 3.**
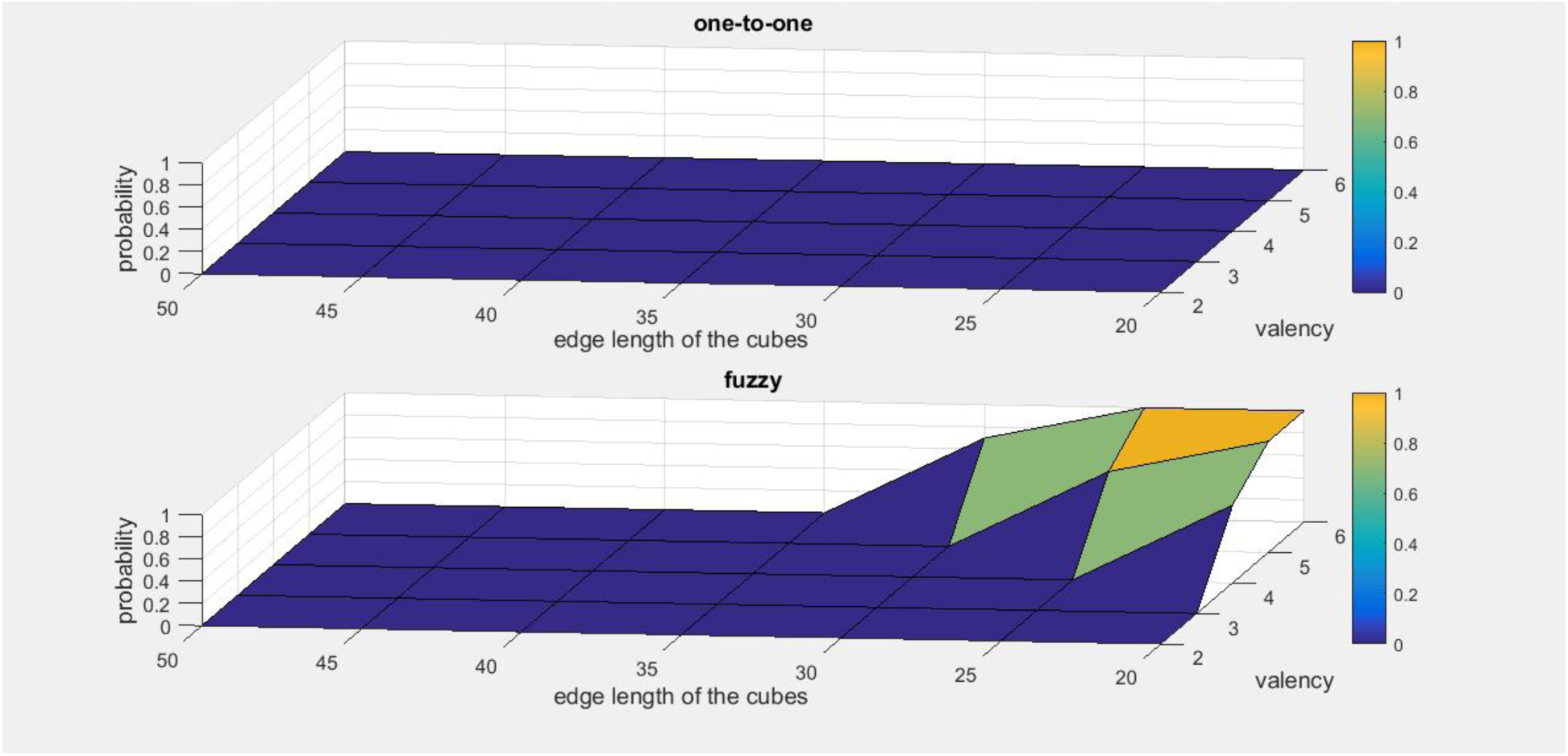
*Probability of large polymers as a function of valency and edge length in one-to-one (**upper**) and fuzzy (**lower**) binding models. Valency is defined as the number of binding elements. Concentration is given by the number of molecules/volume (L^3^). Binding element affinity=0.35, linker dynamics=0.35.*

Higher affinity between the interacting elements increases the association probability (eq. 10) and thus requires lower valency for polymerization as reflected by both one-to-one and fuzzy simulations (Figure 4 upper panel). Binding affinity has a pronounced effect on the one-to-one binding model and values > 0.4 can induce polymer formation with binding elements > 4 (using the same conditions as in Figure 3, where no aggregation was observed). In the fuzzy simulations, polymerization occurs at lower affinity (Figure 4 lower panel), illustrating that partial, heterogeneous binding may compensate for lower affinity in formation of higher-order assemblies. This is consistent with abundance of weakly interacting motifs, such as cation-pi, pi-pi, aromatic hydrogen bonds in proteins composing membraneless organelles [10, 11]. Obviously, above a certain limit, increasing affinity and valency induces formation of aggregates or amyloid structures and not dynamical assemblies, as it will be discussed later.

**Figure 4.**
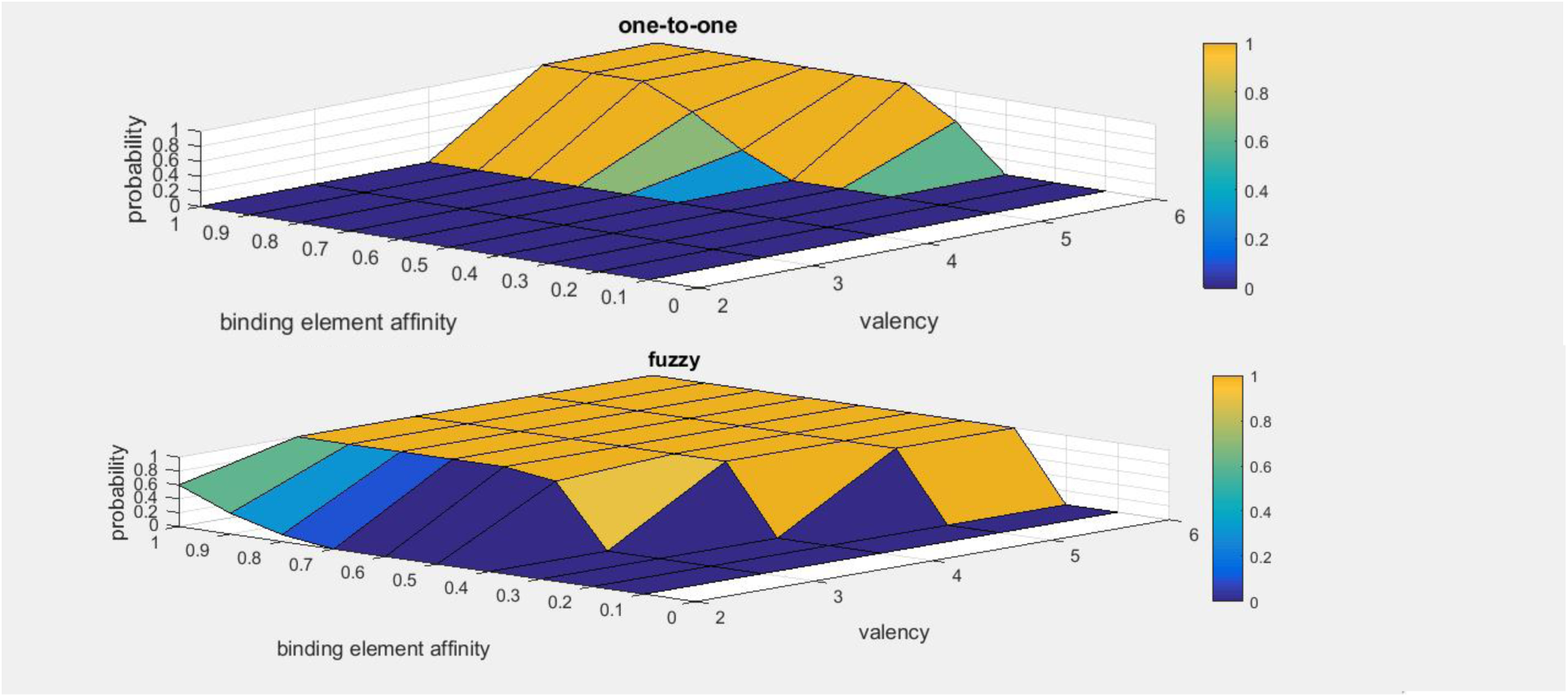
*Probability of large polymers as a function of valency and binding element affinity in one-to-one (**upper**) and fuzzy (**lower**) binding models. Valency is defined as the number of binding elements, which affinity (S_αi_) is computed as the average of the residue-based values (eq. 1). Linker dynamics=0.35, box length = 20.*

The fuzzy model can take the effect of local concentration of the binding elements into account via the modified affinity values (eq. 9), which are incorporated into both association and dissociation probabilities (eq. 10, eq. 11). As anticipated, higher local concentration of the binding sites considerably lowers the phase boundary, even in case of lower valency (Figure 5). The partial, heterogeneous interactions in the fuzzy model correspond to those in an encounter complex [34], which were shown to facilitate productive contacts.

**Figure 5.**
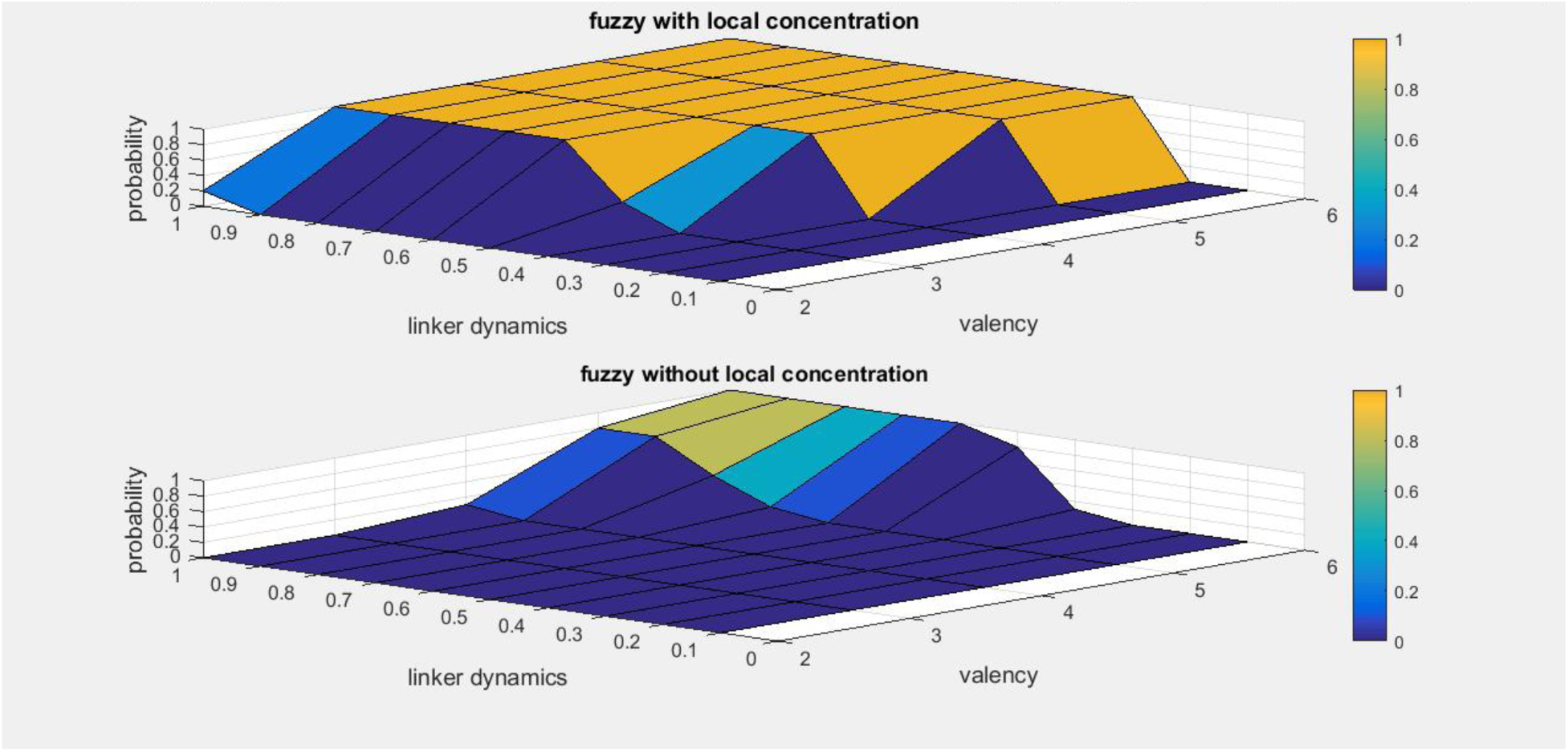
*Probability of large polymers as a function of valency and binding affinity in the fuzzy binding model with (**upper**) and without (**lower**) considering the local concentration effect. Local concentration is computed by eq. 9. Linker dynamics=0.35, box length = 20.*

Intrinsically disordered regions influence phase boundary via increasing linker dynamics [21, 26]. In most studies, linkers are implicitly assumed to have infinite flexibility to enable all possible contact combinations in higher-order assemblies. Limited linker dynamics however, would reduce the number of contact topologies (i.e. microstates) realized in the system. This effect has never been studied systematically. In Figure 6 we present the dependence of polymer formation on linker dynamics, which was varied in the range of [0,1], while keeping the affinity of binding elements constant. If D=1, linkers preserve their conformational heterogeneity similarly to their unbound state [19], while in case of D= 0, the linkers collapse and become rigid in the assembly. The latter case is hypothetical, and is never realized in biological systems.

**Figure 6.**
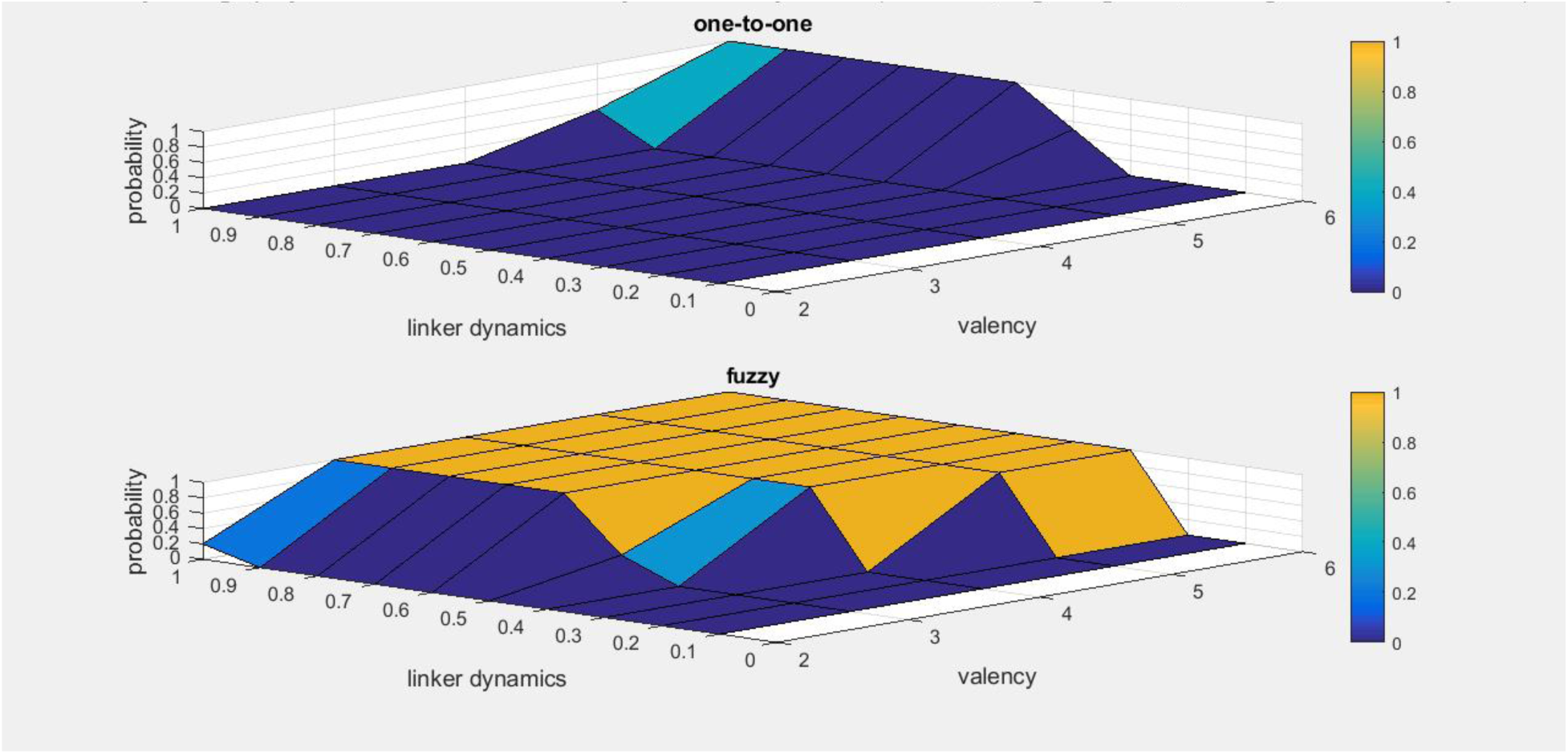
*Probability of large polymers as a function of valency and linker dynamics in the oneto-one (**upper**) fuzzy (**lower**) binding models. Linker dynamics is computed as the average of the residue-based values (eq. 5.). Binding element affinity=0.35, box length = 20.*

Increasing linker dynamics significantly lowers the phase boundary in both one-to-one and fuzzy simulations (Figure 6). In the non-fuzzy binding model, the impact of linker dynamics is comparable to that of increasing interaction affinity (Figure 4), illustrating that either stronger or more heterogeneous binding can promote of assembly formation. Linker dynamics has larger impact on the fuzzy simulations, which demonstrates two effects. First, it highlights that preserving conformational heterogeneity in the assembled state has a critical role in mediating a multitude of binding arrangements. Second, interaction heterogeneity correlates with increasing tendency for phase transition at a given valency.

The interplay between affinity and linker dynamics in the fuzzy simulations is shown in Figures 7-9. Here we systematically varied both quantities in the [0,1] range. For any combinations of binding affinity and linker dynamics, the critical role of valency is observed.

**Figure 7.**
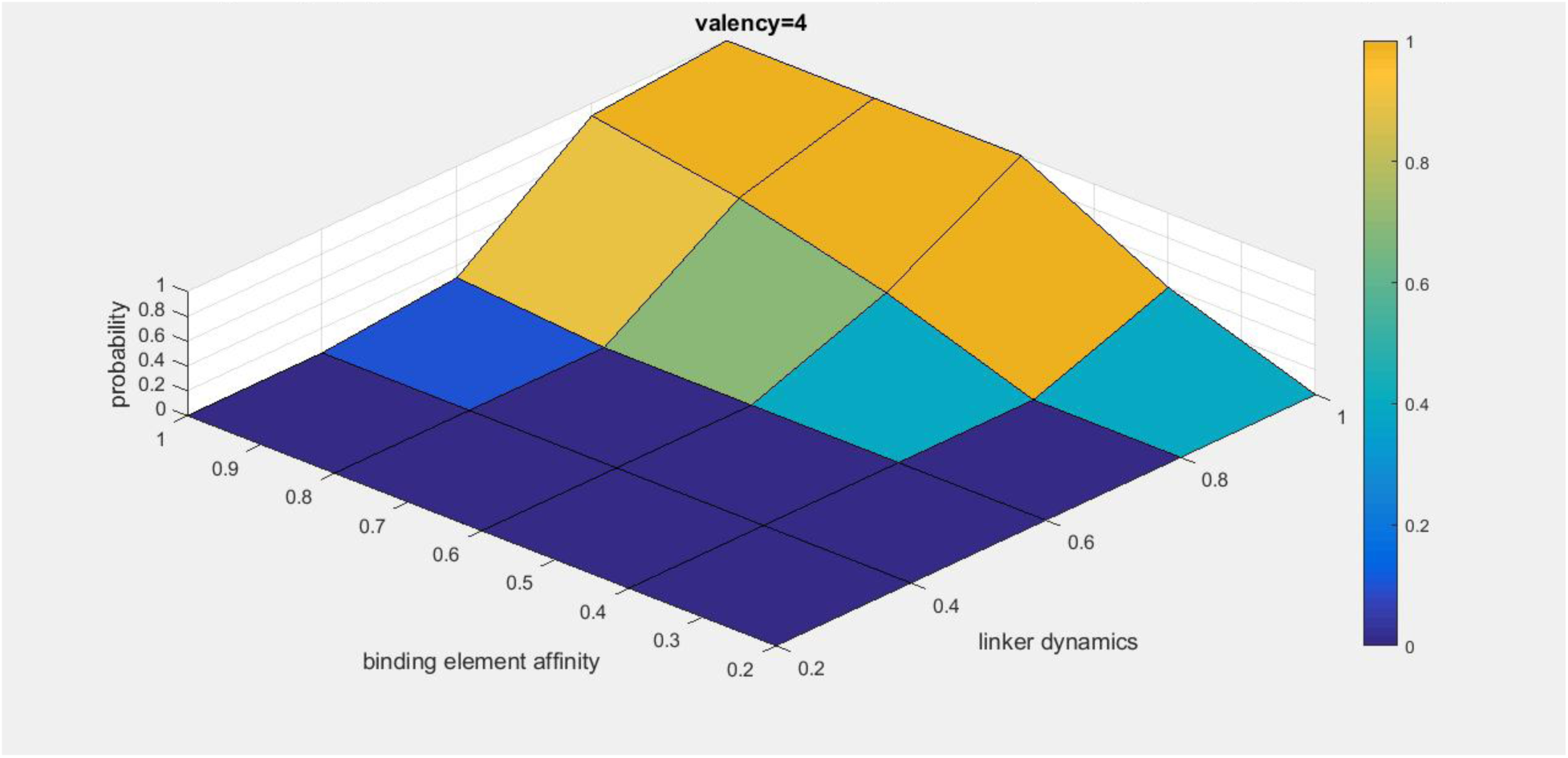
*Probability of large polymers as a function of binding element affinity and linker dynamics in the fuzzy binding models for valency n=4.*

**Figure 8.**
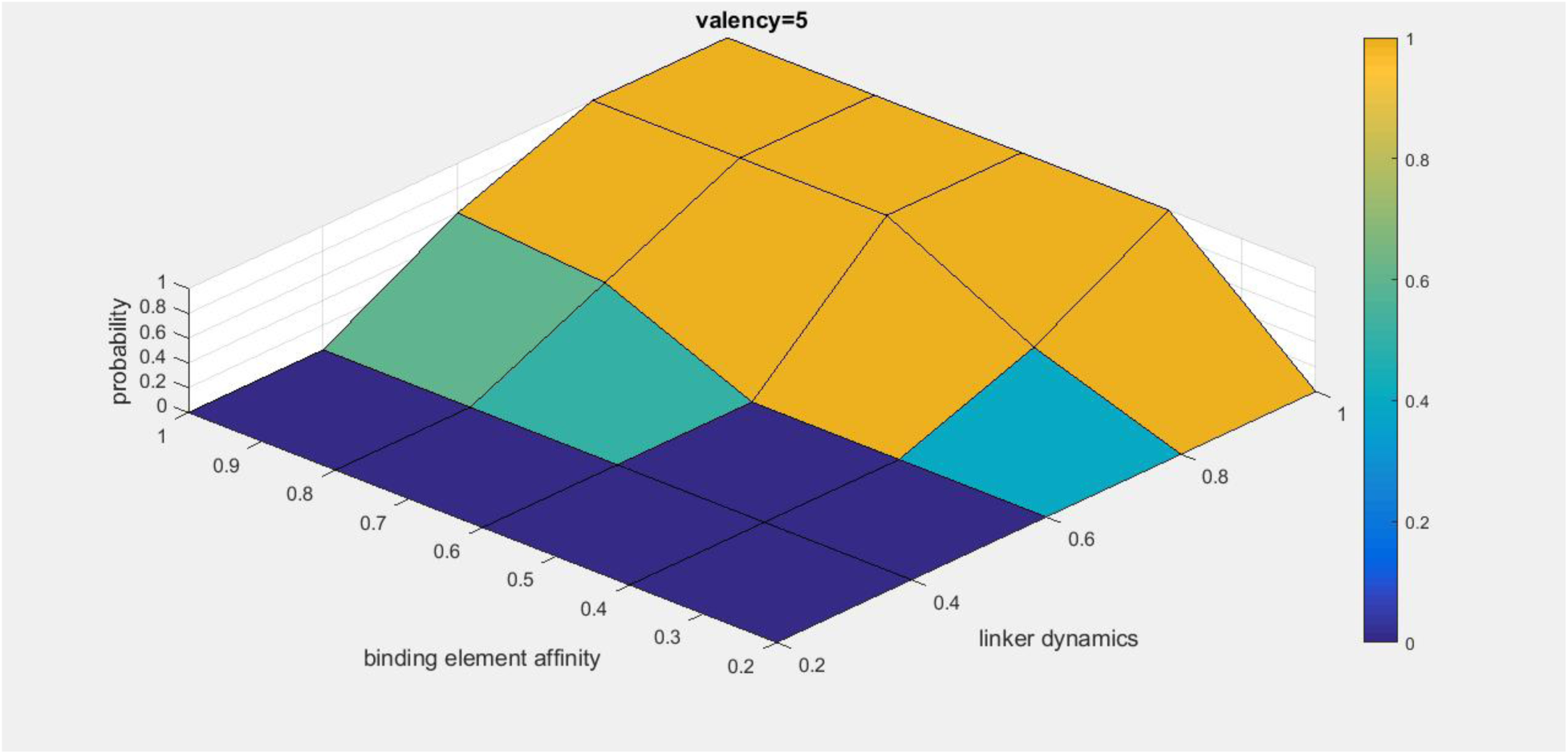
*Probability of large polymers as a function of binding element affinity and linker dynamics in the fuzzy binding models for valency n=5.*

**Figure 9.**
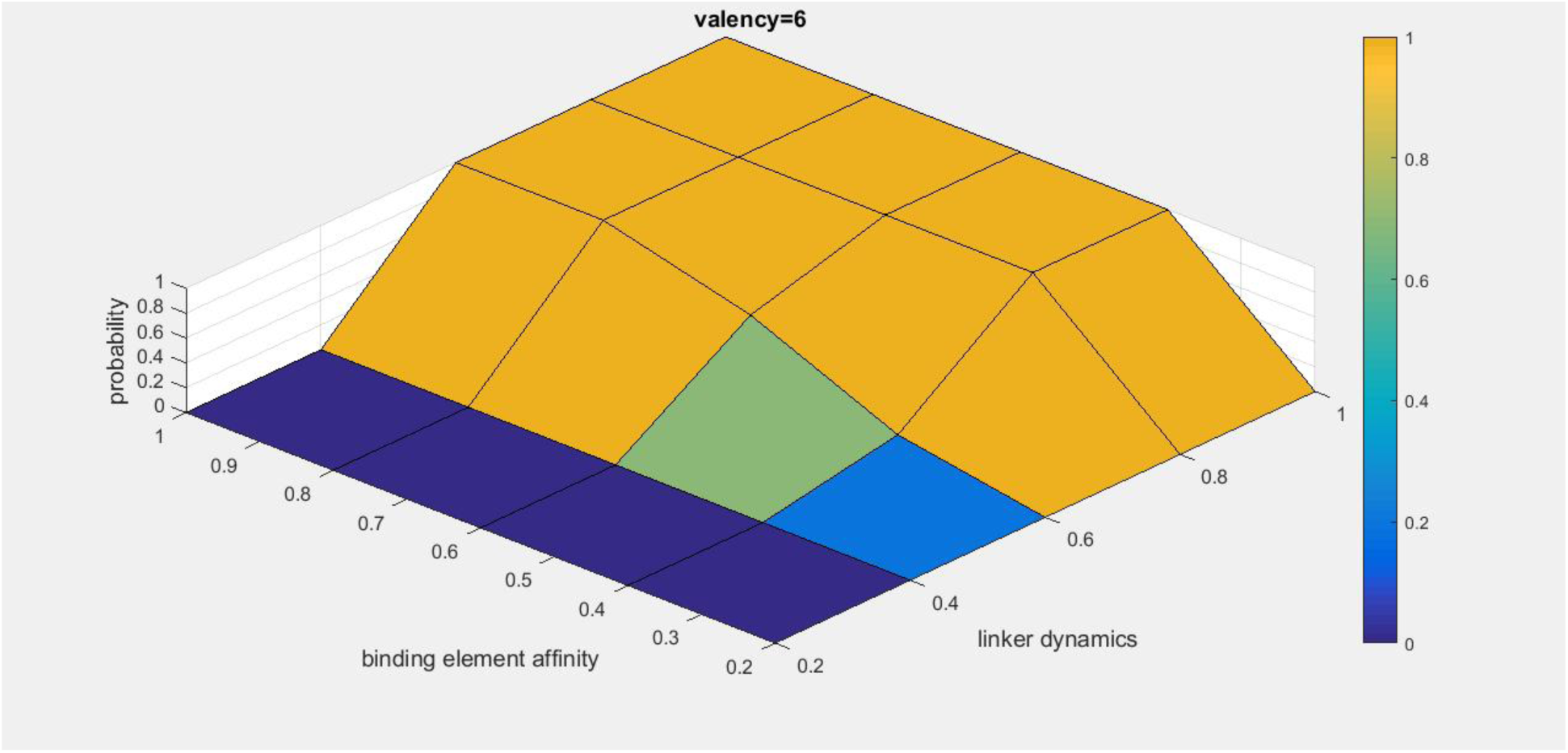
*Probability of large polymers as a function of binding element affinity and linker dynamics in the fuzzy binding models for valency n=6.*

Interaction affinity and linker dynamics act in synergy to promote polymerization. Increasing linker dynamics result in phase transition with lower affinity elements. In turn, higher affinity binding enables polymerization with more rigid linkers. Increasing affinity or dynamics represent two alternative modes for assembly, which result in supramolecular assemblies with distinct properties. Stronger interactions will bias for more ordered, static higher-order assemblies, whereas dynamical linkers lead to structurally heterogeneous, liquid like systems. We must note that our simple model cannot inform on the specificity of interactions, especially in more dynamical systems.

The impact of modulating affinity and linker dynamics on the material state could be evaluated based on the entropy of the system. This was computed according to the reference [14] as

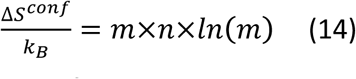

where Δ*S*^*conf*^ is the configurational entropy, *k*_*B*_ is the Boltzmann constant, *n* is the valency and *m* is the number of molecules within the polymer. Figure 10 shows the relative variation of entropy upon systematic changes in binding element affinity and linker dynamics in case of valency n=5.

**Figure 10.**
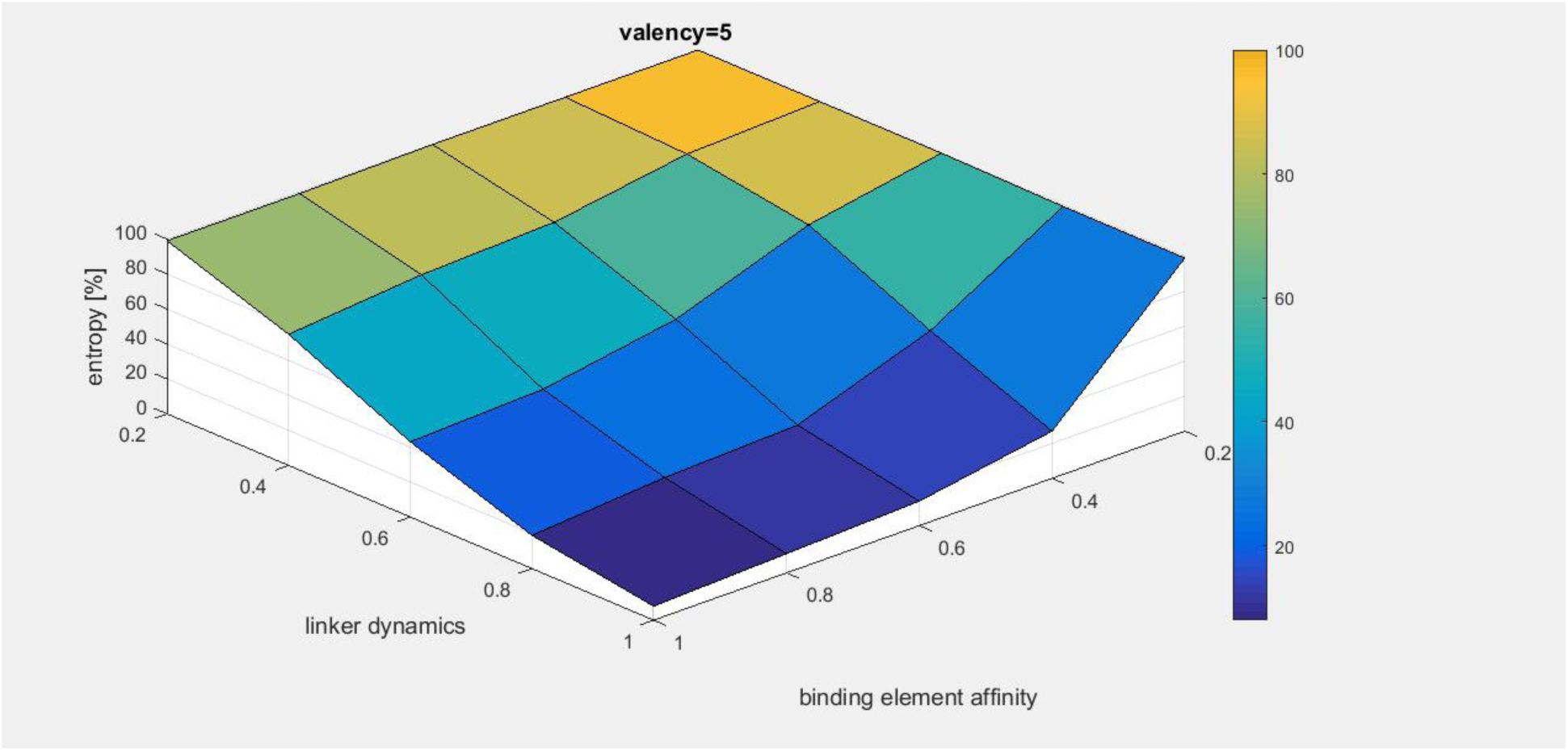
**Relative entropies generated by different strategies of assembly: modulating binding element affinity and linker dynamics.**

The entropy of the system is tuned by the interplay between affinity and linker dynamics. Stronger interactions promote polymerization, especially at higher valency, but freeze the system at the same time. More dynamical linkers may compensate this effect and increase entropy by enabling higher number of microstates. The present simulations do not capture the entropic penalty due to the loss of conformational entropy, which could also be lowered by preserving conformational heterogeneity in the bound system [19].

## Discussion

Protein function is usually interpreted within the deterministic framework of the classical structure-function paradigm. This relationship establishes a connection between a well-defined three-dimensional organization of amino acid residues and the biological activity of the resulted conformer. The classical description also involves the assumption that the intra- or intermolecular interactions generate a well-defined pattern. Increasing experimental evidence contradict this simple picture and demonstrate that biological function may require conformation and interaction heterogeneity [23, 30]. Sequences of proteins composing membraneless organelles for example, are enriched in redundant/degenerate motifs, which appear to contact in multiple ways resulting in a heterogeneous assembly [19]. Indeed, structural and interaction heterogeneity is an intrinsic feature of higher-order protein assemblies, ranging from static to highly dynamical structures [21].

Developing computational approaches to describe heterogeneous systems is a challenge. Until now a one-to-one binding model has been employed in both coarse-grained and lattice simulations [26], which could not account for the effect of heterogeneity, resulted by multiple, alternative configurations. A fuzzy mathematical framework allows co-existing alternative structures or interaction patterns in the system, which are realized to different extents. Within the fuzzy model, one binding element may interact with multiple partners simultaneously and the contribution to alternative states are expressed via membership functions. The membership of a binding element varies in each configuration, and the system remains heterogeneous throughout the trajectory.

Here we applied the fuzzy framework to a hypothetical polymer characterized by binding affinity and dynamics. Although our fuzzy model is highly intuitive, it could describe the basic features of a real protein, especially with a low-complexity sequence. The simulations recapitulate the observation that multi-valency is a pre-requisite for phase transition [14]. As compared to the one-to-one (non-fuzzy) binding model, the fuzzy simulations predict a lower phase boundary (Figure 2,3). This illustrates that higher number of iso-energetic sub-states favor assembly. Furthermore, more partial contacts (local concentration effect) increased the chance of productive interactions and polymerization (Figure 5). Linker dynamics has a distinguished role in increasing heterogeneity, which facilitates formation of large polymers especially in the fuzzy model (Figure 6). We could also systematically investigate the interplay between binding affinity and linker dynamics (Figure 7-9). These two factors present two alternative ways to promote higher-order assembly: higher affinity for the binding elements bias for more ordered states, whereas more mobile linkers increase dynamics of the bound system (Figure 10). Taken together, fuzzy simulations capture the inherent heterogeneity of higher-order protein assemblies leading to an efficient computational technique, which can be applied to studying pathological mutations leading to more solid aggregates.

## Conclusion

Understanding the molecular basis of how higher-order protein assemblies are organized and regulated is challenging owing to the complexity and heterogeneity of these systems. Here we applied a mathematical model based on a fuzzy framework, where biological polymers are described by multiple co-existing states. To our knowledge this is the first time, when such an approach has been applied to protein systems. Fuzzy simulations could more efficiently recapitulate the experimental observations on multivalent polymers as compared to the one-to-one binding model. We propose that the fuzzy framework is generally applicable to real protein systems, not only to the hypothetical model used in this study. The affinity and dynamical parameters could be computed based on the primary sequence. Other parameters could also be incorporated into the fuzzy framework thus opening new perspectives for simulating how complex protein systems work.

## Acknowledgement

We thank Pal Jedlovszky Jr. for his assistance with the simulations and critical reading of the manuscript, and Tibor Sarvari for fruitful discussions. M.F acknowledges financial support of GINOP-2.3.2-15-2016-00044 and the Hungarian Academy of Sciences.

